# Nutritional dependence of sperm mitochondrial metabolism and small RNA biogenesis

**DOI:** 10.1101/2021.10.20.465156

**Authors:** Rashmi Ramesh, Signe Skog, Daniel Nätt, Unn Kugelberg, Lovisa Örkenby, Anita Öst

## Abstract

A wide spectrum of exogenous factors, including diet, environmental pollutants, stress, and seasonal changes have major impact on sperm quality and function. The molecular basis, however, that explains this susceptibility remains largely unknown. Using a combination of proteomics and small RNA (sRNA) sequencing, we show that *Drosophila* sperm display rapid molecular changes in response to dietary sugar, both in terms of metabolic/redox proteins and sRNA content, particularly miRNA and mitochondria derived sRNA (mt-sRNA). Thus, results from two independent omics point at the dynamics of mitochondria as the central aspect in rapid metabolic adjustments in sperm. Using specific stains and *in vivo* redox reporter flies, we show that diet indeed rapidly alters the production of mitochondrial derived reactive oxygen species (ROS). Quenching ROS via supplementation of N acetyl cysteine reduces diet-upregulated miRNA, but not mitochondrial-sRNA. Together, these results open new territories in our search for the mechanistic understanding of sperm health and disease.

**Highlights:**

- Diet rapidly changes the proteomic and sRNA profiles in sperm
- Diet sensitive sperm proteins are found in human infertility studies
- Sperm mitochondrial ROS levels are modulated by diet
- dme-miR-10 regulation is secondary to diet-induced ROS
- Diet, but not diet-induced ROS, alters the expression of mitochondrial small RNA, especially tsRNA

## Introduction

The male germ cell, the harbinger of subsequent generation’s genetic material is surprisingly sensitive to environmental perturbations. In addition to endogenous factors, environmental and lifestyle-related factors including metabolic disorders such as obesity and type II diabetes impacts sperm quality, causing male infertility (Cescon, Chianese, & Tavares, 2020; Chianese et al., 2017; Day et al., 2016; Du Plessis et al., 2010; Katib, 2015; Liu & Ding, 2017; Pergialiotis et al., 2016; Schagdarsurengin & Steger, 2016; Tavares et al., 2016). Such sensitivity of the sperm has been assumed to be secondary to suboptimal spermatogenesis, and consequently, exploratory studies investigating links between environmental shifts and sperm function have employed long-term or chronic intervention. Recently, however, we and others, have found that interventions shorter than the duration of spermatogenesis modulate the molecular composition of the sperm (Gapp et al., 2021; Nätt et al., 2019; Trigg et al., 2021). An enticing and plausible explanation for such rapid responses is that exosomes produced in the epididymis transport material to the sperm during the maturation process (Sharma et al., 2018; Trigg et al., 2021).

Sperm-borne small RNA have been shown to be sensitive to environmental perturbations (Donkin et al., 2016; Fullston et al., 2016; Fullston et al., 2013; Hua et al., 2019; Nätt et al., 2019) and actively influence offspring phenotypes (Chen, Yan, Cao, et al., 2016; Gapp et al., 2014; Grandjean et al., 2015; Rassoulzadegan et al., 2006; Rodgers, Morgan, Leu, & Bale, 2015; Sharma et al., 2016; Trigg et al., 2021). In parallel, links between sRNA and male infertility are becoming increasing clear, with sRNA being suggested as potential biomarkers for male reproductive health (Abu-Halima et al., 2014; Hua et al., 2019; B. Nixon et al., 2019; Salas-Huetos et al., 2016). This duality of sperm sRNA as a potential biomarker for infertility and their importance in early embryonic development, combined with our finding that some diet-sensitive sRNA correlates with sperm motility (Nätt et al., 2019), suggests a shared aetiology for male factor infertility and intergenerational metabolic responses (Nätt & Öst, 2020).

The sperm mitochondria are important for optimal motility and full reproductive potential. Subsequently, mitochondrial function has been studied in relation to male infertility (recently review in Boguenet et al., 2021). For example, comparative proteomics of human sperm have pinpointed mitochondrial proteins to be downregulated in patients with athenozoospermia (Agarwal et al., 2016; Amaral et al., 2014; Bui et al., 2018; Moscatelli et al., 2019; Nowicka-Bauer et al., 2018). Additionally, mitochondria are primary ROS producers in the sperm (Koppers, et al., 2008; Kothari et al., 2010) and ROS acts as a double-edged sword-providing for basal physiologic function of sperm as well as creating oxidative damage. ROS-production has been studied in the context of male infertility (Agarwal et al., 2019; Evans et al., 2021; MacLeod, 1943; Tremellen, 2008). Despite recent advances facilitated by the usage of high-throughput techniques, such as transcriptomics, metabolomics and proteomics providing insights into the basic molecular composition and mechanisms of spermatogenesis and male infertility (Carrell et al., 2016), little is known about the mechanisms connecting rapid responses to diet, molecular changes in sperm and ultimately male fertility. With the rise in global male infertility (Sengupta, Dutta, & Krajewska-Kulak, 2017) there is a need for a comprehensive approach to longitudinally link paternal diet to changes observed in sperm.

Using our *Drosophila* model for paternal intergenerational metabolic response (IGMR) (Öst et al., 2014), we demonstrate that both proteomic and sRNA profiles in sperm are dynamic. We observe that certain diet-regulated proteins are known markers of male infertility, thus strengthening the link between diet and fertility. A short dietary intervention with varying amounts of sugar also resulted in rapid changes in the sperm mitochondrial ROS production. We show that miRNA-especially the upregulation of miR-10 is a result of diet-induced ROS. Next, we focused on tRNA derived fragments (tsRNA), which has been implicated in diet-induced intergenerational responses in many species (Chen et al., 2016; Nätt et al., 2019; Sharma et al., 2016). Finally, combining our highly controlled *Drosophila* model with human sperm sRNA data we identified mitochondrial tsRNA as major components in diet-dependent regulation of sperm across species.

## Results

### A brief dietary intervention rapidly modifies the sperm proteome

We first explored the proteomic landscape of the sperm and sperm microenvironment using proteomics. Briefly, seminal vesicles with mature sperm were dissected for protein extraction and subjected to mass spectrometry (Figure 1A). In total, 542 proteins were identified (Figure 1A-pie chart, Table S1), of which 149 proteins were also found in at least two previously published studies (Dorus et al., 2006; Takemori & Yamamoto, 2009; Wasbrough et al., 2010) (Table S2). Moreover, 65 proteins in our dataset (Table S3), were also present in MitoDrome- a database for nuclear encoded-mitochondrial proteins in *Drosophila melanogaster* (Sardiello, et al., 2003; D’Elia et al., 2006). Importantly, the nine most abundant proteins in our dataset were all sperm-specific (Figure S1A), including sperm-exclusive proteins such as loopin-1 and β-*T*ubulin 85D (betaTub85D) (Bastian et al., 2021). Functionally, most of the identified proteins were involved in cellular metabolic pathways, while others belonged to the broad categories of translation and protein synthesis, heat-shock response, and accessory gland proteins (Figure 1A-pie chart, Table S1). The substantial coverage of our data with published studies, along with the identification of primarily metabolic proteins in sperm, justified further functional investigation in relation to dietary intervention.

**Figure 1:**
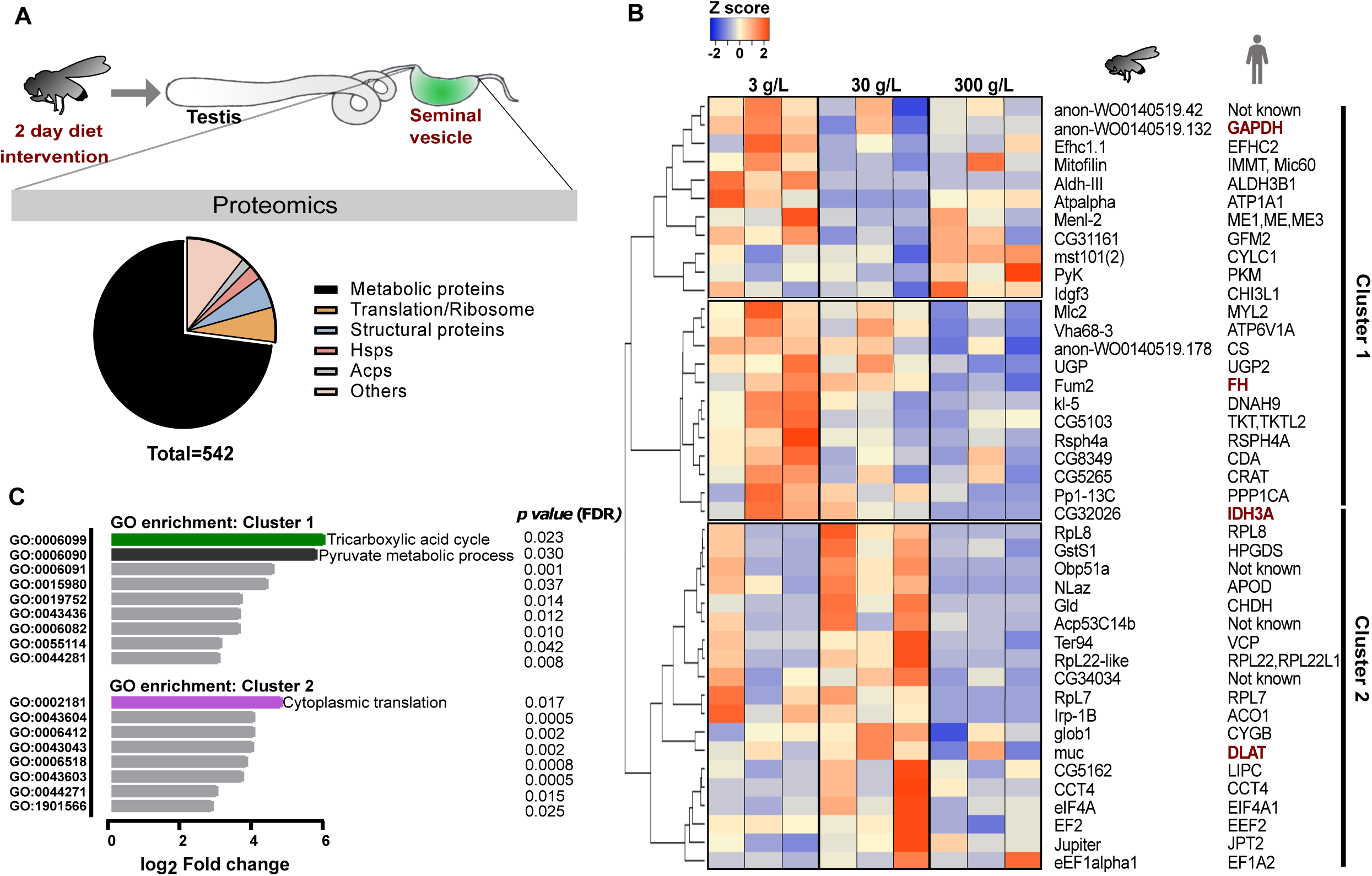
Rapid response to dietary sugar involves proteomic shifts in sperm. **A**: Schematics and Functional distribution of sperm proteome. 2-3-day old virgin *W*^*1118*^ males were fed a diet containing 3-, 30- or 300 g/L sugar for 2 days. Seminal vesicles were dissected out and proteomics was performed by mass spectrometry. Pie chart reveals the proteomic distribution. Metabolic proteins constitute most of the proteome (in black), followed by proteins in other categories (in colour). **B**: Heatmap of significantly changed proteins (*p<0*.*08*). Fly proteins and their human orthologues are shown under their respective cartoon pictograms. N=3, with n=20 per replicate, Pearsons distance based hierarchical clustering with complete linkage-based clustering. Clusters 1 and 2 are indicated with grey bars on the right. Significance test was performed with a linear model fit and Benjamini Hochberg adjustment using R package limma version 3.42.2. All comparisons are made to 30 g/L condition. Protein biomarkers of human male infertility are written in red text. **C:** GO term and statistical overrepresentation analyses of proteins in clusters 1 and 2 using *pantherdb*. Log_2_ Fold change calculated based on the entire fly genome is on x axis, and GO terms are in y axis. Coloured bars indicate the highest fold changes. False discovery rate (*p value*) is given for each GO term. *See also Figure S1*

Three groups of adult virgin male flies were, therefore, fed differing amounts of sugar (10-fold increases viz 3, 30 or 300 g/L) for two days (Figure 1A). A subset of proteins was differentially expressed (Figure 1B), but the most abundant sperm specific proteins - such as loopin-1 and *beta*Tub85D - remained stable in all three dietary conditions (Figure S1B). Unsupervised hierarchical clustering of the differentially expressed proteins (*p*<0.08) revealed two major clusters (Figure 1B, clusters 1 & 2). Whilst proteins involved in TCA cycle (CS, Fum2, CG32026, Irp-1B) and pyruvate metabolism (PyK, Gapdh and Menl-2) were enriched in cluster 1, translation-related proteins were dominating cluster 2 (Rpl22-like, Rpl7, RpL8, eIF4A, EF2) (Figure 1C). Strikingly, proteins in each category (*viz* TCA, pyruvate metabolism or redox) showed a similar trend of expression across the conditions tested (Figure 1B, 2A and 2E), indicating that these metabolic pathways are rapidly modulated by diet in sperm. To gain more translational and functional insights, we studied their human orthologues *in silico*. Interestingly, many of the proteins present in Figure 1B had previously been reported in human male infertility (CTD Gene-Disease Associations dataset) (Rouillard et al., 2016). Among these were well studied protein biomarkers for male infertility such as IDH3A, GAPDHS, FH, DLAT (Figure 1B, red text) (Agarwal et al., 2020; Torra-Massana et al., 2021). This translational overlap (Figure 1B) supports the hypothesis of a link between paternal diet (sugar) and infertility. We, therefore, tested the infertility phenotype in flies via RNAi mediated knockdown of single candidate proteins in the germline (Figure S1C). Among the tested RNAi lines, knockdown of eIF4A, CCT4 and mitofilin resulted in strong phenotypes- with smaller testes and a complete absence of mature sperm (representative micrographs for eIF4A knockdown in Figure S1D).

In summary, our findings highlight the rapid shift in the sperm proteome after a brief dietary intervention. Notably, the human orthologues of the shifted proteome are implicated in male infertility providing important clues about a mechanism associating diet with male infertility.

### Diet acutely modulates ROS production in the sperm

Careful inspection of the differentially expressed proteins (Figure 1B) revealed a subset of proteins that is involved in stress /redox homeostasis (Figure 2A). The expression of these proteins, namely Gsts1, Nlaz, Gld, eIF4A, TER94, glob1 and PP1-13c, were higher in 30 g/L compared to the other two diets (Figure 2A). We, therefore, hypothesised that the flies eating 30 g/L sugar would differ in sperm ROS production compared to flies eating more or less sugar.

**Figure 2:**
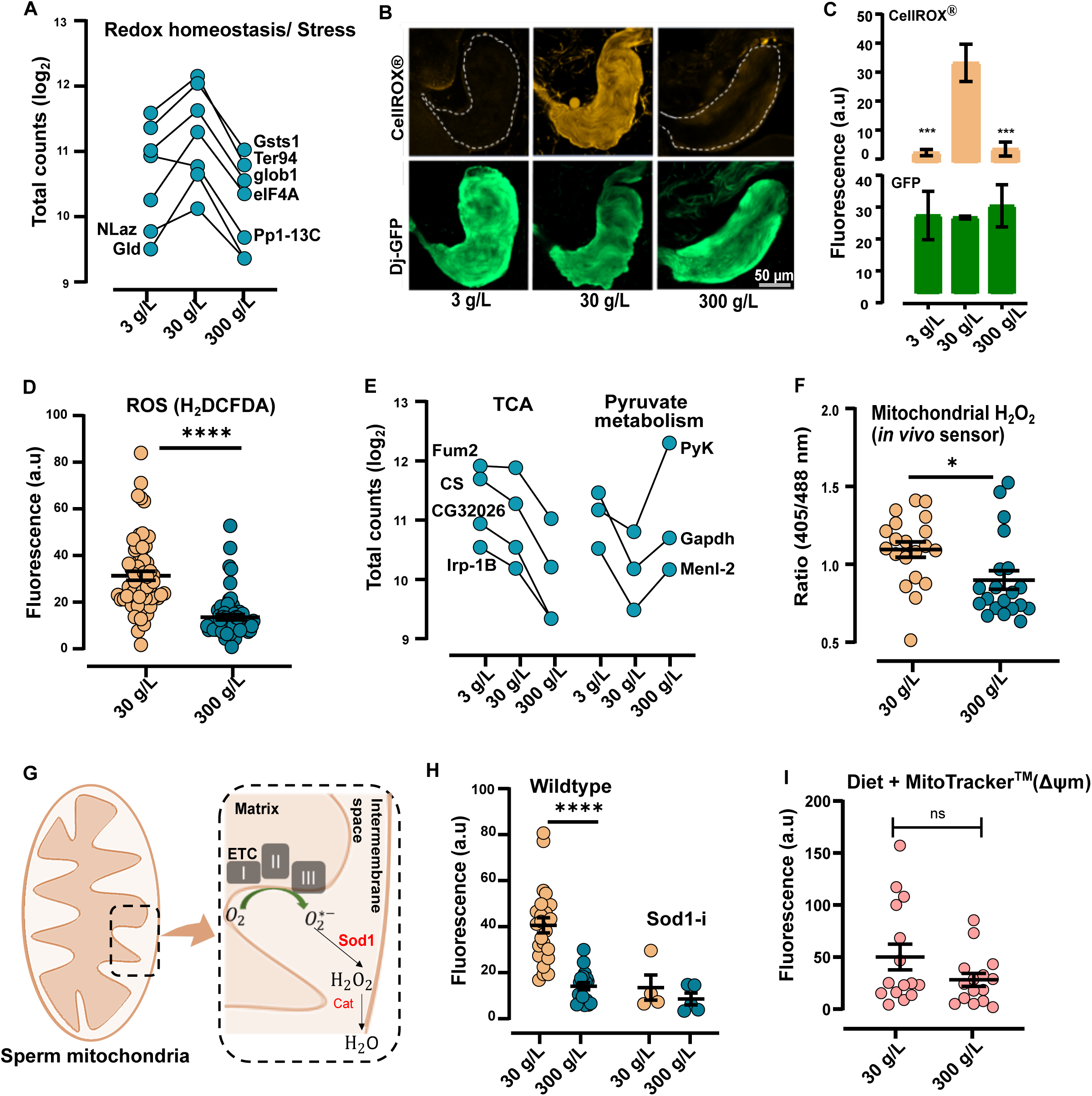
Paternal diet rapidly changes mitochondrial ROS production in sperm. **A:** Proteins involved in stress/ redox homeostasis found in Figure 1B are plotted separately to indicate expression trends across the three diets. Mean Log_2_ levels (total counts) are on y axis, the different sugar concentrations are indicated on x axis. **B:** Visualisation of ROS in seminal vesicle. Flies were fed different diets as in Figure 1A and their seminal vesicles were incubated with CellRox® dye. Images represent ROS in mature sperm in orange channel (CellRox® orange) and Dj-GFP (sperm tail) in green channel. ROS is seen clearly in 30 g/L (middle panel), with minimal to no orange fluorescence seen in 3- or 300 g/L (top and bottom panels). n=4-6, scale bar= 50 µm. **C:** Quantification of ROS labelled with CellROX®. Seminal vesicles from B were quantified using Fiji. CellRox® and GFP channels were separately quantified. Orange bars represent CellRox®, and green bars represent the corresponding GFP quantification. Asterisks (*) represent *p ≤ 0*.*001*, unpaired t-test. **D:** Quantification of ROS labelled with H_2_DCFDA. Flies were fed different diets as in Figure 1A and seminal vesicles were incubated with a different ROS labelling dye-H_2_DCFDA. Fluorescence was quantified using Fiji. n=58-60, asterisks (*) represent *p ≤ 0*.*0001*, unpaired t-test. **E:** Proteins involved in pyruvate metabolism and tricarboxylic acid cycle (TCA) found in Figure 1B are plotted separately to indicate expression trends across the three diets. **F:** Quantification of redox changes in mitochondrial H_2_O_2_ in seminal vesicles of flies expressing the mito-roGFP2-orp1 ratiometric sensor. Ratio of fluorescence between 405 and 488 nm wavelengths are plotted on y axis, and diets are indicated on x axis. n=20, asterisks (*) denote *p ≤ 0*.*05*. unpaired t-test. **G:** Cartoon representation of sperm mitochondria, with the electron transport chain (ETC) zoomed in (dotted box). Complexes I, II and III of the ETC are highlighted, and superoxide production from oxygen is indicated (O_2.-_) in the intermembrane space. Sod1 and catalase enzymes present in the mitochondrial intermembrane space are shown to catalyse sequential reactions converting O_2.-_ to H_2_O_2,_ and H_2_O respectively. **H**: Quantification of ROS levels in wildtype flies and in flies with germline specific knockdown via RNAi of Sod1 (Sod1-i). n=4-5 (Sod1-i) and n=18-24 (Wildtype), unpaired t-test. **I**: Quantification of mitochondrial potential (ΔΨm). Fluorescence intensities from seminal vesicles stained with Mitotracker™ CMXRos. n=15, unpaired t-test. **Note:** All comparisons were made to 30 g/L condition (except A and E). The data shown are mean ± SEM. All quantification are done using Fiji (in C,D,F,H,I) and mean grey values are plotted on y-axis and diet is indicated on x axis (except F). a.u=arbitrary units, ns=no significant *See also Figure S2*

Seminal vesicles from flies with a sperm-specific GFP-reporter -*Don juan* GFP (*Dj*-GFP), that ate 3-, 30- or 300 g/L sugar for two days, were dissected and incubated with the ROS-indicator (CellROX® Orange). Visualization of the live tissues revealed a striking difference in ROS production in response to diet (Figure 2B). Sperm from 30 g/L sugar diet showed distinct orange fluorescence indicative of active ROS production, while negligible/no fluorescence was detected in low- and high-sugar diets (Figure 2B, C-upper panel). The differences in fluorescence were not a result of changes in the amount of sperm *per se*, since GFP fluorescence corresponding to *Dj*-GFP from sperm tails remained unchanged (Figure 2B, 2C; lower panel). We also independently replicated the findings in a separate fly strain (W^*1118*^) using a general ROS indicator dye (H_2_DCFDA) (Figure 2D). Together, this clearly suggests that a diet comprising 30 g/L sugar promotes ROS production in the sperm of *Drosophila melanogaster*.

Proteins involved in TCA cycle were upregulated in seminal vesicles of 30 g/L sugar-eating males (Figure1B, 2E), whilst proteins involved in glycolysis were downregulated (Figure1B, 2E). Given that the TCA cycle operates in the mitochondria, and that mitochondria are significant ROS producers in sperm (Koppers et al., 2008; Kothari et al., 2010) we hypothesised that diet-induced ROS originates in the mitochondria.

We, therefore, used a previously characterised ratiometric H_2_O_2_ redox sensor, mitochondrial-roGFP2-Orp1 (Albrecht et al, 2011) (Figure S2A-C). This *in vivo* sensor allowed us to specifically track mitochondrial ROS, with the added advantage of defining H_2_O_2_ as the specific ROS species modulated by diet. As previously described (Figure 1A), 3–5-day old male flies were fed 30- or 300 g/L sugar for 2 days, and fluorescence was measured by confocal microscopy by sequential excitation at 405- and 488 nm (See methods for details). In line with the TCA-related protein changes, we found that 30 g/L condition presented seminal vesicles with a higher 405/488 ratio suggesting the involvement of the mitochondria in creating the oxidative environment via H_2_O_2_ production (Figure 2F, Figure S2C).

In mitochondria, superoxide radicals are produced mainly from the electron transport chain (ETC) and are converted to H_2_O_2_ by the enzyme superoxide dehydrogenase (SOD) (Figure 2G). To validate that diet modulates the mitochondrial H_2_O_2_ production, we initiated a germline specific knockdown of the mitochondrial enzyme-Sod1 (Figure S1C) and measured ROS levels by microscopy (as in Figure 2B, C). As expected, quantification of fluorescence in seminal vesicles after RNAi revealed a reduction in the amounts of mitochondrial ROS irrespective of diet, while the genetically unmodified controls on a 30 g/L sugar diet showed ROS generation (Figure 2H). Together, these results highlight the prominent redox changes in sperm mitochondria in response to diet.

We next turned to investigating whether mitochondrial function itself was modulated by diet. Since 30 g/L favoured ROS production in comparison to 300 g/L sugar, we reasoned that either mitochondrial morphology or activity was altered. We, therefore, measured mitochondrial potential (ΔΨm). Seminal vesicles from flies eating 30- or 300 g/L sugar for two days were dissected and incubated with a fluorescent dye that specifically stains the mitochondria (MitoTracker™ Red CMRos). The uptake of this cell-permeable dye is dependent on ΔΨm, whereby a strong signal of the dye indicates a high mitochondrial potential. Bright red fluorescence was seen in seminal vesicles from flies feeding either 30- or 300 g/L sugar (Figure S2D). Quantification of red fluorescence in seminal vesicles (containing mature sperm) revealed no significant differences in sperm of either diet (Figure 2I). ROS levels in seminal vesicles were measured like in Figure 2D on a separate set of flies from the same experiment, and as observed before, 30 g/L favoured ROS production (Figure S2E). These results suggest that although diet had a rapid effect on ROS production in sperm, there was little to no effect on the mitochondrial ΔΨm in mature sperm. Although a general change in ΔΨm was not detected, it is possible that specific sites in the ETC are affected by diet. Indeed, previous studies where individual ETC complexes were modulated, support ROS-mediated signalling via both conventional as well as reverse electron transport (reviewed in (Scialò, Fernández-Ayala, & Sanz, 2017)). Thus, future studies on dissecting the role of individual complexes are needed. What is clear, however, is that a dietary change of just two days is reflected in the sperm mitochondrial H_2_O_2_ production.

### Acute dietary intervention rapidly modifies the sRNA profiles in sperm

Having found that mitochondrial ROS respond rapidly to diet, we sought to explore whether sperm sRNA profiles were similarly shifted, and if so, whether such changes would be secondary to diet-induced ROS. For this, we took advantage of the direct quenching of ROS with general antioxidants such as N acetyl cysteine (NAC) supplemented in diet containing either 30- or 300 g/L sugar (Figure 3A). After two days, the seminal vesicles were dissected and incubated with CellROX®, and ROS was measured by microscopy and quantified (Figure 3A). As previously observed, the 30 g/L sugar diet resulted in high ROS production in sperm, while flies fed 300 g/L sugar had little to no ROS (Figure 3A, -NAC). NAC supplementation (1 mg/mL) in both 30- and 300 g/L diets greatly diminished the ROS levels (Figure 3A, +NAC), indicating the effectiveness of the antioxidant in quenching the diet-induced ROS in seminal vesicles. We therefore used NAC to investigate the effect of diet-induced ROS on sperm sRNA. Importantly, to eliminate the contribution from somatic sRNA, only pure mature sperm was used for sRNA sequencing (Figure 3A, B). Sperm sRNA sequencing revealed the presence of different sRNA biotypes including, in order of their abundance, rRNA-derived small RNA (rsRNA), tRNA-derived small RNA (tsRNA), mitochondrial tRNA-derived small RNA (mt tsRNA), microRNA (miRNA), long intergenic non-coding RNA (lincRNA), piwi-interacting RNAs (piRNA) and sRNA from protein-coding RNA (Table S4). For comparison, we analysed human sperm sRNA data from Nätt et al. 2019. Reads mapping to rsRNA dominate the sperm of both flies and human (94% versus 74% respectively, see Table S4 for rsRNA information in fly sperm). We did not detect gross changes in the nuclear rsRNA profiles and have therefore excluded them from our analyses due to their high presence.

**Figure 3:**
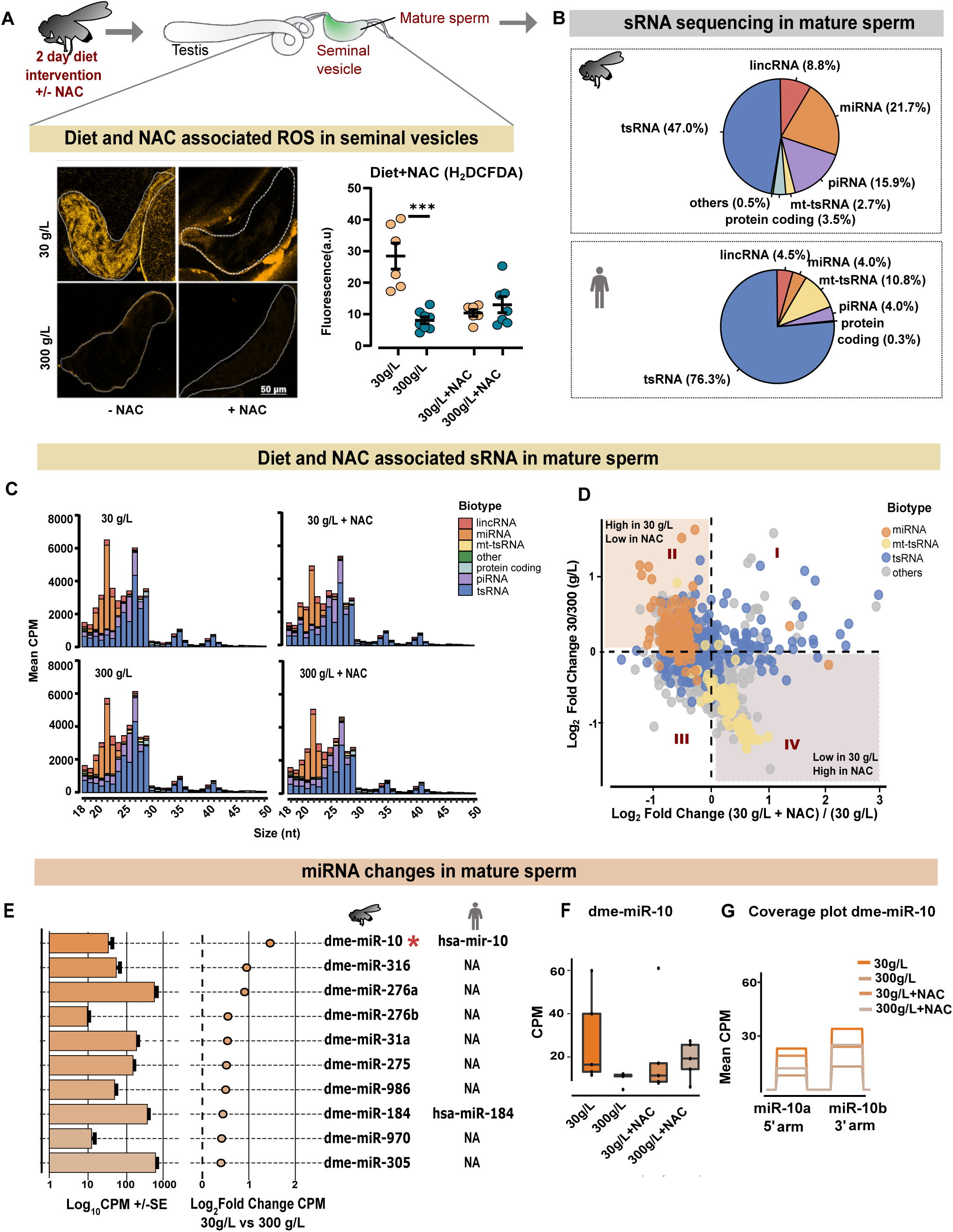
Rapid alteration in sperm sRNA following dietary intervention. **A:** Addition of antioxidant N-Acetyl Cysteine (NAC) mitigates ROS levels in sperm. Following a 2-day dietary intervention, seminal vesicles were dissected out and ROS levels were visualised using H_2_DCFDA and quantified. The data shown are mean ± SEM. On a separate set of flies from the same experiment, sperm was isolated and sequenced for sRNA. **B:** Comparison of sRNA in fly and human sperm. Human sperm sRNA data were derived from (Nätt et al., 2019). Shown in pie charts are percent CPM of perfect matches to respective reference genome. Reads mapping to rsRNA are not shown. Reads mapping to snoRNA (less than 2%), snRNA (less than 1%), and reads with no annotation to any of the sRNA-subtype (less than 1%) are classified as “others”. Fly data n=20, human data n=15. **C:** Size distribution of sRNA in *Drosophila* sperm in the diets with and without NAC. Reads mapping to rsRNA are excluded. Reads mapping to snoRNA, snRNA and those sRNA that map to the reference genome but not to any other sRNA-subtypes are consolidated as “other”. **D:** Scatter plot of fold changes showing the distribution of sRNA in the tested conditions. x axis represents Log_2_ fold change (30 g/L+NAC) / (30 g/L) and y axis represents Log_2_ fold change (30 g/L) / (300 g/L). Log fold changes was calculated on CPM. Each point represents one individual sequence, in total 983. Coloured closed circles are shown for miRNA (orange), mitochondrial tRNA (yellow), tRNA (blue) and others (grey). Reads mapping to rsRNA are not included. **E:** miRNA identified in this study. Fly miRNA are indicated under the fly pictogram. Bars represent mean CPM. The corresponding closed circles represent mean Log_2_ fold change. Error bars are ± SE. Human orthologues, where present, are indicated under the human pictogram. Asterisks (*) indicate significance (*p<0*.*05*) calculated by a Generalized Linear Negative Binomial model for all sequences that originate from the same miRNA. NA= Not Available. **F:** Bar graph for CPM expression of dme-miR-10 in all four dietary groups. Each point represents the mean value of the two miRNA sequences mapping to dme-miR-10 for each sample. **G:** Coverage plot of dme-miR-10 showing expression of both the 5’ (dme-miR-10a) and 3’ arms (dme-miR-10b). Each coloured line represents mean CPM of indicated dietary condition. CPM=counts per million *See also Figure S3*

A side-by-side comparison of sRNA from flies and human sperm (Nätt et al. 2019) revealed striking similarities in the distribution and abundance of various sRNA biotypes, with tsRNA being the abundant biotype in sperm of both species (Figure 3B), as previously reported in mature sperm in mice (Peng et al., 2012). The abundance of the other sRNA biotypes varied more between fly and human sperm. Notably, compared to human, miRNA and piRNA had a stronger presence in fly sperm (Figure 3B). Nonetheless, global similarities of sRNA biotypes between fly and human justifies the use of a fly model to better understand mechanisms in human sperm.

Next, we looked at the size distribution of various sRNA biotypes in sperm of flies fed with 30- and 300 g/L sugar with and without NAC (Figure 3C). The size distribution showed diverse lengths of tsRNA, spanning 18-50 nucleotides (Figure 3C). Notably, the proportions of the various biotypes in the range of 18-30 nucleotides remained similar across the four conditions, suggesting that diet or NAC did not induce a major shift in the profiles of these sRNA (Figure 3C). The fold change distribution of sRNA biotypes, however, revealed more specific effects between conditions (Figure 3D). Whilst tsRNA showed a diverse spread across the conditions (Figure 3D, closed blue circles), miRNA were predominantly upregulated in 30 g/L condition (Figure 3D, quadrant II, and Figure S3A, B). In parallel, mt-tsRNA were downregulated in the 30 g/L condition (Figure 3D, closed yellow circles, quadrant IV, and Figure S3A).

Previously, we have shown that a high sugar diet in humans resulted in the upregulation of several miRNAs (Nätt et al., 2019). In the current study, almost all miRNA were upregulated in 30 g/L diet both in the presence and absence of NAC. Intrigued by the finding that most miRNAs (84.8% and 98.3% respectively, Figure S3 A, B & Table S5) were upregulated, we decided to further investigate specific miRNAs. Interestingly, the most differentially expressed miRNA -dme-miR-10 (CPM of 30 g/L vs 300 g/L) reached clear statistical significance in the 30 g/L condition (*p<0*.*05*, Figure 3E asterisk *, Figure 3F). It has been shown that the rodent and human orthologue, miR-10, is diet responsive (Cropley et al., 2016), and is upregulated in male infertility of both rodents and human (Gao et al., 2019). Among the upregulated miRNA (Figure 3E & Figure S3C), is miR-276b, which is a known regulator of synchronous egg-hatching in locusts (He et al., 2016), suggesting that sperm miRNA may have long-lasting effects in early embryogenesis. Interestingly, in the presence of NAC, miR-10 is downregulated (Figure 3F, miR-10). Both the 5’ and 3’ arms of miR-10, called miR-10a and miR-10b respectively, were upregulated in the 30 g/L sperm and similarly quenched by NAC (Figure 3G), making miR-10 a strong candidate for future research. The upregulation of miRNA in the ROS-producing diet (30 g/L sugar), and the mitigation of this effect by NAC supports the idea that miRNA regulation is secondary to the diet-induced ROS production in sperm.

### Sperm tsRNA show varied responses to diet and ROS

We next analysed the tsRNAs in both 30- and 300 g/L sugar diets with and without NAC (Figure 4, Table S6). First, all transcripts mapping to full-length tRNAs were classified based on cut-sites using the bioinformatic package-Seqpac (Skog et al., 2021) and with information about tRNA loop structure taken from tRNAscan-SE (Lowe & Chan, 2016). Five sub-types of tsRNA were defined: 5′-half, 5′-tsRNA, i’-tsRNA, 3’-tsRNA and 3′-half (Figure 4A). The distributions of the tsRNA subtypes varied depending on whether they were of nuclear or mitochondrial origin (Figure 4B), suggesting alternative pathways of biogenesis. The nuclear tsRNA were dominated by 5’-halves (Figure 4B-nuclear tsRNA, & Figure S4A), as reported previously (Chen et al., 2016; Nätt et al., 2019; Sharma et al., 2016). On the other hand, i’-tsRNA were predominant amongst the mitochondrial tsRNA (Figure 4B-mitochondrial tsRNA). This indicates distinct cleavage signatures in the two sub-cellular compartments. More importantly, the tRNA cleavage sites were mainly dependent on the transcriptional origin of the tRNA rather than the diet itself (Figure S4B, C).

**Figure 4:**
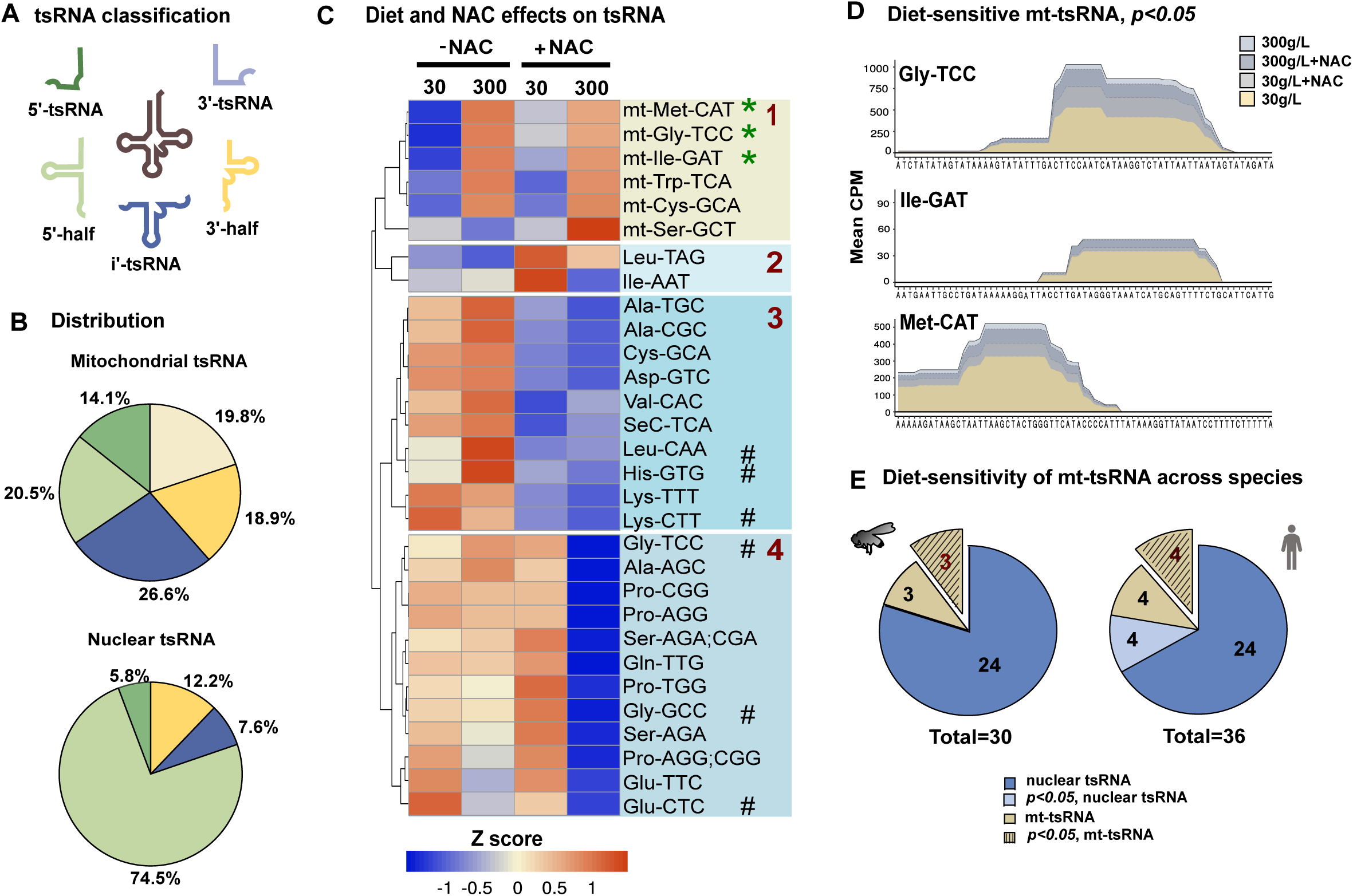
Sperm tsRNA have varied responses to diet and ROS. **A:** Illustrative representations of the fragments derived from tRNA analysed in this study. See Methods section for details on tsRNA classification **B:** Distribution of tsRNA based on their mapping to nuclear and mitochondrial genomes. For each tsRNA, percentage of mean CPM is presented. **C:** Heatmap showing expression of tsRNA. TsRNAs that originate from the same mature tRNA are combined. Each column represents the mean CPM expression from the indicated dietary condition. Clusters are based on unsupervised clustering and are numbered 1-4 in red. The ‘mt’ in cluster 1 indicates tsRNA of mitochondrial origin. Asterisk (*) represents statistical significance (*p<0*.*05*) as determined by Negative Binomial Generalized Linear Model. Hashtag (#) refers to tsRNA identified in other studies (Chen et al., 2016, Sharma et al., 2016). **D**: Coverage plots of significantly altered mitochondrial tsRNAs (from C), namely Gly-TCC, Ile-GAT and Met-CAT. x axis shows the corresponding mature tRNA sequence, and y axis represents the mean CPM values. **E**: Dietary regulation of mitochondrial tsRNAs is conserved from fly to man. Mt-tsRNA are represented in yellow plain pies, with longitudinal black lined yellow pies representing changes reaching statistically significance. Blue pies represent the nuclear tsRNAs. In both flies (this study) and humans (Nätt et al., 2019), half of all reported mitochondrial tsRNAs were significantly different with sugar diet. Statistical significance (*p<0*.*05*) was determined by Negative Binomial Generalized Linear Model. *See also Figure S4*

The observed downregulation of mt-tsRNA in the 30 g/L condition (Figure 3D), and the tsRNA cleavage signature in the mitochondria (Figure 4B) raised an intriguing possibility that tsRNA biogenesis in the mitochondria might be a direct consequence of diet-mediated responses. Therefore, to get an overview on ROS mediated tRNA cleavage, all fragments carrying the same anticodon sequence (isodecoders and isoacceptors) were combined and mean CPM values were visualized as a heatmap (Figure 4C). Unsupervised clustering of these tsRNA revealed four clusters (Figure 4C, clusters 1-4). Interestingly, the mitochondrial tsRNA (Figure 4C, highlighted in blue text) and nuclear tsRNA (Figure 4C, black text) formed separate clusters, as previously observed in human sperm (Nätt et al., 2019), which again indicates different biogenesis pathways. The mitochondrial tsRNA were generally less abundant in 30 g/L compared to 300 g/L diet (Figure 4C, 30-& 300 g/L, -NAC). This downregulation of some mitochondrial tsRNA in 30 g/L, namely Ile-GAT, Gly-TCC and Met-CAT was statistically significant (Figure 4C, asterisks* & Figure 4D). Most importantly, the presence of NAC did not alter the diet-induced shift of mitochondrial tsRNA (Figure 4C, 30-& 300 g/L, +NAC). This supports the hypothesis that changes in mt-tsRNA happens prior to diet-induced mitochondrial ROS.

The nuclear pool of tsRNA (Figure 4C, clusters 2-4) included fragments previously identified as being diet-sensitive (Figure 4C, see #) (Chen et al., 2016; Sharma et al., 2016). Notably, three sub-clusters emerged among the nuclear tsRNA (Figure 4C, clusters 2-4). TsRNA in cluster 3 had a higher presence in both diets (Figure 4C, 30-& 300 g/L, -NAC), while addition of NAC reduced the levels of the same tsRNA (Figure 4C, 30-& 300 g/L, +NAC), indicating alternative sources of tRNA fragmentation in the presence of NAC. It is plausible that non-diet induced ROS, could have been quenched by the presence of NAC, giving rise to these fragments. The nuclear tsRNAs in cluster 4 show indistinct effects of both diet and NAC (Figure 4C, cluster 4). This suggests that multiple mechanisms are at play and ROS or diet, alone or in combination exert multiple effects on certain tsRNA.

Together, these findings point at dynamic changes in the tsRNA profiles in sperm mitochondria in response to diet, further highlighting the role of mitochondria in diet-induced metabolic alterations in sperm. In fact, comparing diet-sensitive mt-tsRNA in humans (Nätt et al., 2019) and flies shows striking similarities, with at least 50% of the observed mt-tsRNA being altered by diet (Figure 4E).

## Discussion

Here, we have used proteomics and sRNA sequencing, combined with in-depth bioinformatic analyses in the *Drosophila* model of paternal intergenerational metabolic response (IGMR), to identify the molecular changes in the sperm. A short dietary intervention with varying amounts of sugar resulted in a rapid remodelling of the proteome in seminal vesicles containing mature sperm. Whereas the bulk of structural proteins was unaffected by diet (Figure S1B), we found changes in proteins involved in metabolism and stress /redox homeostasis (Figure 1A, B & Figure 2A). This change in redox homeostasis was reflected in ROS production in sperm mitochondria, which dramatically increased with 30 g/L sugar (Figure 2 B-D, F, & Figure S2C). In parallel, high-throughput sequencing of purified sperm RNA revealed coordinated changes in the sperm sRNA. In depth analyses of the sRNA profiles indicated that response to diet in sperm manifested in different ways. NAC mediated depletion of ROS in sperm was able to reverse the expression of diet-altered miRNA, in particular miR-10 (Figure 3D-E), suggesting this change to be secondary to ROS. The nuclear and mitochondrial tRNA fragments (tsRNA) were also altered and separated into unique clusters suggesting a mixed response to diet and ROS (Figure 4C). More specifically, and in line with findings in human sperm, we found that diet, but not diet-induced ROS, altered the expression of the mt-tsRNAs (Figure 4C, D & Figure S4B, C). Together, our results favour a model where mitochondrial metabolic flexibility and sRNA biogenesis are in the centre of diet-dependent molecular changes in sperm.

Since sperm rely on substrates from their microenvironment to fuel their metabolism, it is easy to envision that temporal changes in nutrient flux are directly mirrored in the sperm metabolism. Indeed, the existence of a gut-gonad axis has been demonstrated in *Drosophila*, wherein the male intestine secretes citrate to the adjacent testes and promotes sperm maturation (Hudry et al., 2019). A similar gut-gonad axis was recently described in a sheep model of diet-induced metabolic syndromes (Zhang et al., 2021). In both these models, metabolic perturbations altered spermatogenesis and sperm numbers. However, with our short dietary intervention, two findings indicate unchanged sperm numbers: (1) highly expressed sperm proteins such as loopin-1 and beta-tubulin are maintained at the same level in the proteomics data independent of diet (Figure S1A), and (2) microscopy of seminal vesicles of the sperm-specific fusion protein Dj-GFP revealed no change in fluorescence intensity (Figure 2C). In addition, staining of seminal vesicles with MitoTracker™ Red CMXRos showed similar staining patterns across the tested diets, revealing negligible to no changes in sperm numbers or in mitochondria potential. It is, however, possible that long term dietary changes, or other nutrient compositions, would impede spermatogenesis and modulate the number of sperm being produced.

While sperm numbers appeared unchanged by diet, we observed that diet altered the mitochondrial H_2_O_2_ production. Given that ROS is required for sperm functions in humans (Du Plessis et al., 2015; Dutta et al., 2020), modulation of such essential processes by diet is intriguing and can have far-reaching consequences. Given its easy diffusibility across membranes, and a longer half-life, H_2_O_2_ is considered one of the main signalling molecules amongst the ROS species (Holmström & Finkel, 2014; Rhee, 2006; Veal, Day, & Morgan, 2007). On the other hand, ROS-induced oxidative stress can lead to male infertility (Barati, Nikzad, & Karimian, 2020;Agarwal, Makker, & Sharma, 2008; Agarwal et al., 2019; Bui et al., 2018; Tremellen, 2008). Therefore, production of ROS must be counterbalanced by antioxidant systems. In mammals, it is well known that proteins secreted from the epididymis provide such a protection for the maturing sperm (reviewed in Chianese & Pierantoni, 2021). Similarly, we find a rapid upregulation of certain enzymes involved in redox homeostasis – such as Gsts1 and Glob1 in the high ROS condition (Figure 2A). Considering the transcriptional and translational quiescence of the sperm, it is likely that somatic cells in the seminal vesicle has a similar role as the epididymis in providing an antioxidant environment for the sperm. In mammals, proteins (Candenas & Chianese, 2020; Nixon et al., 2019) and sRNA made in somatic cells are known to be packaged and transported to sperm via extracellular vesicles (Belleannée et al., 2013; Sharma et al., 2016; Vojtech et al., 2014; Xu et al., 2020). For example, miR-10 expression have been shown high in seminal exosome in humans (Barceló et al., 2018; Vojtech et al., 2014), and in mice tsRNA, especially the tsRNA 5’-halves, are known to be loaded onto the sperm via exosomes originating from the epididymis (Sharma et al., 2016; Stanger et al., 2020). Although no changes in the abundance of tsRNA 5’-halves were detected in this study, the markers of extracellular vesicles such as Ter94, APOD, GAPDH (Figure 1B) and miR-10 (Figure 3F), were altered with diet, suggesting the involvement of exosomes in creating an antioxidant response in *Drosophila* seminal vesicle.

Alongside the upregulation of miRNA and proteins involved in stress /redox homeostasis interpreted as part of an antioxidant response to increased ROS, our data point to a separate mechanism for the biogenesis of mt-sRNA detected in sperm (Figure S3A, B). In general, very little is known about sRNA originating from the mitochondria, both regarding their biogenesis and their functionality. It has been reported that mitochondria sRNA, in particular mt-piRNA, is abundant in male mice germ cells (Larriba et al., 2018) . It is interesting to note that we observed about 92% of mt-piRNA to be downregulated in the 30 g/L condition (Figure S3A) but we have yet to explore if they have a functional role in sperm and/or in the fertilized egg. Likewise, it remains an open question whether mt-tsRNA carries a similar regulatory function as has been demonstrated with nuclear tsRNA. Like nuclear tsRNA, the mitochondrial tsRNA in fly (Figure 4D) and human (Nätt et al., 2019) sperm have distinct cut-sites, indicating that sperm mt-tsRNA have specific functions. The role of tsRNA in relaying the paternal nutritional status to the offspring has been shown in rodent models (Chen et al., 2016; Sharma et al., 2016). Thus, it is tempting to speculate that mt-tsRNA have similar roles. Nonetheless, we show that cleavage of mt-tsRNA differs from that of nuclear tsRNA (Figure 4B). This supports recent data showing that the biogenesis of nuclear and mitochondrial tsRNAs differs in flies (Molla-Herman et al., 2020).

Sperm sRNA profiles, especially alterations in miRNA have been studied in infertile men (Abu-Halima et al., 2014; Lian et al., 2009; Muñoz, Mata, Bassas, & Larriba, 2015; Salas-Huetos et al., 2016b; C. Wang et al., 2011), and have been suggested as biomarkers of male infertility (Kotaja, 2014; Barbu et al., 2021; Kiani, Salehi, & Mogheiseh, 2019; Salas-Huetos et al., 2020). Additionally, sperm tsRNA, miRNA and rsRNA has recently been shown to correlate with embryo quality (Grosso et al., 2021; Hua et al., 2019; Nätt & Öst, 2020; Xu et al., 2020) and piRNA are involved in paternal diet induced intergenerational response (Lempradl et al., 2021). We have earlier described that a short high-sugar intervention in healthy young men synchronously increases tsRNA and rsRNA coming from the mitochondrial genome and that the increase of mt-tsRNA is positively associated with simultaneous changes in sperm motility (Nätt et al., 2019). Since mitochondrial energy metabolism is intimately linked with motility, this is an intriguing finding. Moreover, reanalysing data from Donkin et. al we observed that obesity is associated with less rsRNA derived from mitochondrial DNA (Donkin et al., 2016; Nätt et al., 2019). In all, our data adds to the findings that sRNA in sperm can be a good biomarker of male reproductive health (Abu-Halima et al., 2014; Hua et al., 2019; B. Nixon et al., 2019; Salas-Huetos et al., 2016) and provides evidence that both the nutrient state as well as the ROS-production of the sperm can influence sperm health.

We conclude that *Drosophila* sperm are susceptible to dietary changes and have identified candidates that could influence metabolic responses in offspring. With the rise in male factor infertility identification of such biomarkers in sperm should therefore be of prime interest in future investigation.

## Acknowledgements

This research was supported by grants from The Swedish Research Council (2015-03141), Ragnar Söderberg’s foundation and, Knut and Alice Wallenberg foundation (2015.0165) to A.Ö. Technical support from staff at core facilities at Linköping university, especially Vesa Loitto (microscopy unit), Annette Molbaek and ñsa Schippert (RNA sequencing) and Maria Turkina (mass spectrometry) is gratefully acknowledged. The authors are grateful to Marie Roth for technical help with dissections, and Anna Asratian and Cecilia Jönsson for comments on the manuscript.

## Author contributions

Conceptualization: A.Ö. and R.R.; Methodology: R.R, S.S, U.K, D.N, L.Ö, A.Ö. Data curation: S.S. Formal analysis, R.R, S.S, D.N, A.Ö., Resources: A.Ö, Software: S.S, D.N., Visualization: R.R, S.S, D.N, A.Ö, Writing – original draft: R.R, S.S, A.Ö, Writing – review & editing: R.R, S.S, U.K, D.N, L.Ö, A.Ö

Funding acquisition: A.Ö Supervision

All authors have approved the final version of this paper.

## Declaration of interests

The authors declare no competing interests.

## Materials and Methods

### Fly stock maintenance and sugar diet administration

A standard laboratory strain W^1118^ was used for proteomics, sRNA sequencing and certain microscopic experiments in this study. Dj-GFP males were used for ROS experiments to visualize sperm tail in seminal vesicles. The flies were inbred for several generations and maintained on standard cornmeal/molasses media at 26 °C. Male flies were isolated within 2 days of eclosure and aged for an additional 2-3 days before switching them to paternal diet intervention food containing 3, 30 or 300 g/L white sugar, for two days.

Standard food: Agar 10 g/l, yeast 28 g/l, cornmeal 68 g/l, molasses 68 g/l, Nipagin 1.5 g/l, propionic acid 5.5 ml/l.

Paternal diet intervention food: Agar 12 g/l, yeast 10 g/l, propionic acid 4,5 ml/l, soy flour 30 g/l and white sugar as indicated.

### Isolation of seminal vesicles and preparation of protein extracts

Seminal vesicles from 20 males for each diet were isolated in insect medium (Sigma # T3160) using fine forceps and collected in 25µL ice cold mili Q water in a 1.5 mL microfuge tube stored on ice. After dissections of the complete set, tissue was lysed mechanically using a fitted pestle (VWR #431-0094), followed by 2 minutes (40 oscillation per second) on a bead shaker (Qiagen Tissue lyser)-centrifuged at 1000g for 5 minutes at 4 °C to remove debris, and supernatant was used for further processing. 20µL of the supernatant was first alkylated in the presence of 10 mM DTT in ammonium bicarbonate (25mM) for 1 hour at 56°C. Alkylation of cysteines was performed using 55mM iodoacetamide prepared in ammonium bicarbonate solution (25 mM) for 1 hour at room temperature in the dark. Following this, proteins were precipitated using ice cold acetone overnight at –20°C. Subsequently centrifuged at 15000 g for 10 minutes in a cooled rotor, and supernatant was aspirated out. The pellet was used for Trypsin digestion (0.005 µg/µL, Pierce #90057) at 37°C overnight. The next morning, after a further boost of trypsin (0.0025 µg/µL) for 3 hours at 37°C, the digested peptides were vacuum-dried, and stored at –2°C. Peptides were resuspended in 12 µL 0.1% formic acid and used in duplicate (5 µL) for mass-spectrometry analysis.

### Mass spectrometry analysis

The peptides were introduced into an LTQ Orbitrap (Thermo, San Jose, CA) mass spectrometer and all MS/MS samples were analysed using Sequest (Thermo Fisher Scientific, San Jose, CA, USA; version IseNode in Proteome Discoverer 1.4.0.288). Sequest was set up to search Fly uniprot 7227.fasta assuming the digestion enzyme trypsin, with the following parameter settings: 1 miscleavages, variable methionine oxidation and phosphorylation on serine and threonine, carboxymethyl cysteine as fixed modification, with a fragment ion mass tolerance of 0.50 Da and a parent ion tolerance of 6.0 PPM. Results were merged using Scaffold (Proteome Software) version 3.00.04.

### Criteria for protein identification

Scaffold (version Scaffold_4.10.0, Proteome Software Inc., Portland, OR) was used to validate MS/MS based peptide and protein identifications. Peptide identifications were accepted if they could be established at greater than 95,0 % probability by the Scaffold Local FDR algorithm. Protein identifications were accepted if they could be established at greater than 99,0 % probability and contained at least 2 identified peptides. Protein probabilities were assigned by the Protein Prophet algorithm (Nesvizhskii et al; 2003). Proteins that contained similar peptides and could not be differentiated based on MS/MS analysis alone were grouped to satisfy the principles of parsimony. Proteins sharing significant peptide evidence were grouped into clusters. The final list of 542 proteins is provided in Table S1.

### Protein classification

Flybase IDs of the entire list (542 proteins) were analysed in FlyMine. The tool for pathway enrichment was used, with normalization to gene length, and Benjamini-Hochberg correction factor of maximum *p* value 0.05. Most proteins were not assigned to any pathways. In such cases, information from Uniprot and FlyBase were used to assign a broad category for the protein. Most metabolic proteins were assigned by FlyMine. Many proteins were involved in more than one pathway, as apparent in the Table S1. Certain proteins, for example Gld and Glob1, although known to be involved in redox homeostasis, were not annotated by FlyMine. In such cases they were assigned to a class manually. The human orthologues are from FlyBase/DIOPT.

### Bioinformatic analysis (proteomics)

The peptide counts for each protein across the diets tested were compiled into a excel file with individual proteins represented in rows, and each sample per diet in columns. Normalisation to mean of sum of all protein counts was performed. Statistical analyses were performed in R Ver. 3.6.3. Significant changes were analysed with a linear model fit in limma ver 3.42.2 (Ritchie et al., 2015) and edgeR ver. 3.28.1 (McCarthy, Chen, & Smyth, 2012) and adjusted with Benjamini Hochberg for adjusted *p values*. 30 g/L was used as intercept in design. Heatmaps were generated using *heatmapper* (Babicki et al., 2016) hierarchical clustering based on Pearson complete distance. All other statistics were done using GraphPad prism (8.3.0).

### GO term analyses

GO terms for biological processes were assigned using PantherDB (Mi et al., 2021) with the statistical enrichment tool. Gene names from each cluster (Figure 1D) were separately analysed, and the list was sorted based on fold enrichment with reference to the entire genome of *Drosophila*. The fold enrichment was converted to log_2_ values and plotted with GO terms as a bar graph using GraphPad prism (8.3.0).

### Measurement of ROS by microscopy

After paternal diet intervention for 2 days, seminal vesicles were dissected (from Dj-GFP or W^1118^) in insect medium (Sigma # T3160) (for CellROX®) or 1X PBS (H_2_DCFDA), and subsequently incubated in CellROX® orange dye (Invitrogen # C10443) for 30 minutes at 37°C (5 µM final concentration in insect medium) or H_2_DCFDA (Invitrogen #D399) at room temperature for 5 minutes (40 µM final concentration). Following this, the tissue was rinsed extensively with 1X PBS and mounted onto glass slides with halocarbon oil (HC700, Sigma) as the mounting medium, and covered with glass coverslips. Coverslips were sealed with clear nail polish and slides were imaged using an inverted confocal microscope (LSM800, Zeiss) using absorption/emission maxima of ∼545/565 nm (for CellROX®) or ∼488/517 nm (for H_2_DCFDA and GFP). All images were quantified in Fiji (Schindelin et al., 2012). The steps followed to process the images in Fiji are depicted below.

*>>Fiji*

*>>image import* .*czi*

*>>process>subtract background>rolling ball radius 100 pixels*

*>>Image>adjust>threshold-set threshold min and max: 30-35 to 255*

*>>choose ROI*

*>>measure: Integrated density and mean gray values*

### Antioxidant supplementation to counteract ROS

*N* acetyl cysteine (Sigma # A7250) solubilized in water was used in food containing 30- or 300-g/L sugar at a final concentration of 1 mg/mL, and intervention was carried out for two days at 26°C. Following this, testes were dissected, and seminal vesicles were imaged for ROS production using CellROX® orange dye as described above. Simultaneously, sperm from 15 flies from each condition was isolated and used for sRNA sequencing as described below.

### RNAi crosses

RNAi was initiated in the male germline by crossing males expressing UAS-RNAi (SOD1: BDSC#29389; eIF4A: BDSC#33970; CCT4: BDSC#77358) or UAS-GFP RNAi (BDSC#35786) with virgin Nanos Gal4; UAS Dicer 2 (BDSC#25751) on standard food. Eclosed F1 males were subjected to dietary intervention and ROS measurements using CellROX® orange dye as described above.

### Imaging for redox analysis

Flies expressing roGFP2-orp1 in the mitochondria (BDSC# 67672) were subjected to dietary intervention as detailed previously. Flies were dissected in 1X PBS and testes were mounted on halocarbon oil and sealed with coverslips. Imaging was performed immediately using confocal microscopy (upright LSM 700, Zeiss). The probe fluorescence was excited at 405nm and 488nm, sequentially and line by line with emission wavelengths ranging between 518-580 nm. To determine the dynamic range, testes from the 30 g/L sugar diet condition were either fully oxidised (using 10% H_2_O_2_) or fully reduced (1mM DTT) and immediately imaged with the above settings.

### Quantification of redox changes

Fiji was used to quantify each channel (405 nm and 488 nm) separately, with background subtraction and thresholding as described before. To calculate the fluorescent intensity ratios, the 405nm value was divided by 488nm. All ratios were computed in Excel. The dynamic range (DR), which reflects the maximal achievable redox changes in our model, was calculated by dividing the 405/488 ratio of the fully oxidized H_2_O_2_ sample with the same ratio of the fully reduced DTT sample.

### Staining of testes with MitoTracker™ Red CMXRos

Testes of 1- to 3-day-old males were dissected in PBS and stained in MitoTracker™ Red CMXRos in PBS (Molecular Probes, Eugene, OR, USA) for 15 minutes at room temperature (final concentration 1 μM). Tissues were rinsed in 1XPBS and fixed in 4% paraformaldehyde solution for 10 minutes at room temperatures. Following fixation, the tissues were extensively rinsed in 1X PBS triton (0.2%) and mounted on glass slides with VECTASHIELD® containing DAPI (Vectorlabs, H-1200-10). Coverslips were sealed and imaging was performed in an inverted LSM 800 confocal microscope with Texas red filter settings (ex/em: 579/599 nm). Quantification was done after background subtraction and thresholding in Fiji as described before.

### Isolation of sperm for small RNA sequencing

Sperm was isolated in TC-100 Insect Medium (Sigma# T3160) essentially as described in (Öst et al., 2014). From each diet, sperm from 15 flies were dissected and pooled in 1:10 dilution of RNAse inhibitor (Recombinant ribonuclease inhibitor 5 000 U, Cat. 2313A Takara), and samples were flash-frozen on dry-ice, and later stored at -80 °C. For sperm collection, five samples of each diet were prepared.

### Small RNA Library preparation and sequencing

RNA extraction was done using miRNeasy Micro kit (Qiagen, 217084) according to manufactors instructions. Prior to homogenization cold steel beads (0,15 g, SSB02-RNA NextAdvance, Troy NY) were added to frozen samples followed by the addition of 500 ul of prechilled Qiazol (Qiagen). Samples were run in Tissue Lyser LT (Qigen) for 2 + 2 min at 40 oscillations/second.

RNA quailty was studied with BioAnalyzer RNA analysis (5067-1511, Agilent, RNA 6000 nano kit). Small RNA libraries were produced with NEBNext Multiplex SmallRNA Library Prep Kit for Illumina (E7560S, E7580, New England Biolabs) with the customisation of a dilution of primers 1:2. The 3′adaptor ligation reaction was carried out at 4°C overnight. To minimize the amount of 2S rRNA we added a blocking oligo (5′-TAC AAC CCT CAA CCA TAT GTA GTC CAA GCA-SpcC3 3′) to the samples at the 5′ adaptor ligation step ((Wickersheim & Blumenstiel, 2013). Libraries were amplified for 16 cycles and cleaned using Agencourt AMPure XP (Beckman Coulter, Brea, CA). Size selection on amplified libraries was done using TBE gel (EC6265BOX, Invitrogen) 130-165 nt length. Extraction of cDNA from gel was performed with Gel Breaker Tubes (3388-100, IST Engineering) by incubation with buffer included in the NEBNext kit and incubated on a shaker for 1 hour at 37 degrees Celsius, flash frozen for 15 minutes and again incubated on shaker. Gel debris was removed by Spin-X 0,45 µm centrifuge tubes (Corning Inc.,Corning, NY). Percipitation was done using GlycoBlue (Invitrogen), 0,1 times the volume of Acetate 3M (pH5.5), and 3 times the volume of 100% Ethanol in -70°C overnight. Quality of cDNA libraries were studied with BioAnalyzer DNA analysis (5067-1504, Agilent, Agilent High Sensitivity DNA kit, 5067-4626). Final DNA concentration was determined with Quantus Fluorometer (E6150, Promega, Madison, WI) using QuantiFluor ONE ds DNA system. Libraries were pooled and sequenced on NextSeq 500 with NextSeq 500/550 High Output kit version 2.5, 75 cycles (Illumina, San Diego, CA). All libraries passed Illumina′s default quality control.

### Bioinformatic analyses (sRNA)

Data analysis was performed with Seqpac ver. 0.99.0 (Skog et al., 2021). Adaptor trimming, quality control and mapping were all performed in Seqpac with make_counts and make_reanno workflow with an evidence of that an individual sequence should have at least 1 count in 2 separate samples. Trimming was performed on the adaptor sequence of the used NebNext library (AGATCGGAAGAGCACACGTCTGAACTCCAGTCA). Only reads with an adaptor sequence present prior to trimming were studied. Averaged over all 20 samples, 1.9*10^6^ unique reads passed filtering and with a mean of 1,5*10^7^ reads per sample.

Genomic mapping was performed towards *Drosophila* reference dm6 downloaded from UCSC. Biotype mapping was performed to Ensembl ncRNA BDGP6.32, piRNA piRBase D. Melanogaster 2.0 (J. Wang et al., 2019; Yuan et al., 2016; P. Zhang et al., 2014, 2015), miRBase 21 (Griffiths-Jones, 2004; Griffiths-Jones, Grocock, van Dongen, Bateman, & Enright, 2006; Griffiths-Jones, Saini, Van Dongen, & Enright, 2008; Kozomara, Birgaoanu, & Griffiths-Jones, 2019; Kozomara & Griffiths-Jones, 2011, 2014), protein coding from Ensembl BDGP6.32 (Howe et al., 2021)and tRNA from GtRNAdb (Chan & Lowe, 2009, 2016) Drosophila BDGP dm6. Human mapping was performed against the human genome GRCh38.p13 (GCA_000001405.28) from Ensembl. Biotype mapping was performed to Ensembl ncRNA, piRNA piRBase v 2.0, miRBase 21, protein coding from NCBI RefSeq proteins and tRNA from GtRNAdb. Biotypes were hierarchically determined in the order rRNA, mitochondrial tRNA, tRNA, miRNA, snoRNA, lncRNA, snRNA, piRNA and protein coding. Drosophila mitochondrial genome NC_024511.2 was downloaded from NCBI. Bowtie indexes were created with Rbowtie ver 1.32.0.

Data was first filtered with function PAC_filter on a size of 18-50 nt length, 10 counts in 60% of samples and a perfect (no mismatch) match with reference genome. Additional filtering was performed after CPM calculations to remove reads without a presence of minimum 20 counts per million in 25% of samples. Computation of counts per million, log fold change and Figure 3 B, C, E, F and Figure 4 B and D were generated with Seqpac. Other figures presented were created with ggplot2 ver 3.3.3 and pheatmap ver 1.0.12. Unless stated otherwise, data used in figures are CPM for individual sequences. In figure 3 E, log_2_ fold change is calculated on a feature-base rather than sequence base, where all sequences mapping to a certain miRNA are classified together.

Analysis of tsRNA and their cleavage sites were performed with the PAC_mapper and PAC_trna analytic workflow in Seqpac. Here, ss-files constructed with tRNAscan-SE for the Drosophila nuclear and mitochondrial tRNA were used (Lowe & Chan, 2016). We defined five tsRNA subtypes; 5’-half, 5’-tsRNA, i-tsRNA, 3’-tsRNA and 3’-half, where a 5’-half starts in the 5’ end of the mature tRNA and ends in the antiocodon loop. Furthermore, 5’-tsRNA also starts in the 5’ end but ends prior to the antiocodon loop. The opposite relationship is true for 3’-halves and 3’-tsRNA, whilst i-tsRNA are fragments without connection to either 5’ nor 3’ end. Fragments from tRNA are here reported to the isodecoder and isoacceptor they originate from, as most of nuclear tsRNA maps to several copies at once. This multimapping is found in Table S4.

For all source code used, see https://github.com/signeskog/Ramesh-2021.

### Statistical analyses (sRNA)

Statistics on sRNA data were performed with a Negative Binomial Generalized Linear Model, since the data are counts based. sRNA sequences are in some cases impossible to map to one unique place in the genome, due to their short size and the repetitive nature of some transcription sites. As of now, there is no perfect method to add this uncertainty into a statistical model and we have not accounted for the ambiguity stemming from the risk of multimapping. Since we cannot guarantee that each sequence stems from one original place, we did not perform multiple testing.

In the case of miRNAs, we did not take miRNA isoforms into account, but rather combined sequences originating from the same miRNA. This made it so our model studied the difference on a miRNA-to-miRNA basis rather than sequence to sequence basis. The model was performed with function glm.nb from the R package MASS version 7.3-51.4. Clustering of tsRNA (Figure 4 C) were performed with pheatmap v 1.0.12, where clustering was performed on rows with k=4 and with Euclidean distance. Standard error of the mean for log_2_ fold changes were calculated on the SEM of each individual of 30 g/L (n=5) fold change against 300 g/L for each sequence.

## Resource availability

### Lead contact

Further questions may be sent to lead contact Anita Öst.

### Materials availability

All *Drosophila* strains used in this manuscript are commercially available.

### Data and code availability

Raw sRNA-seq performed in sperm have been deposited at Sequence Read Archive (SRA) at accession number PRJNA770968. All code for this project is available on GitHub at https://github.com/signeskog/Ramesh-2021.

## Supplementary information

**Figure S1:**
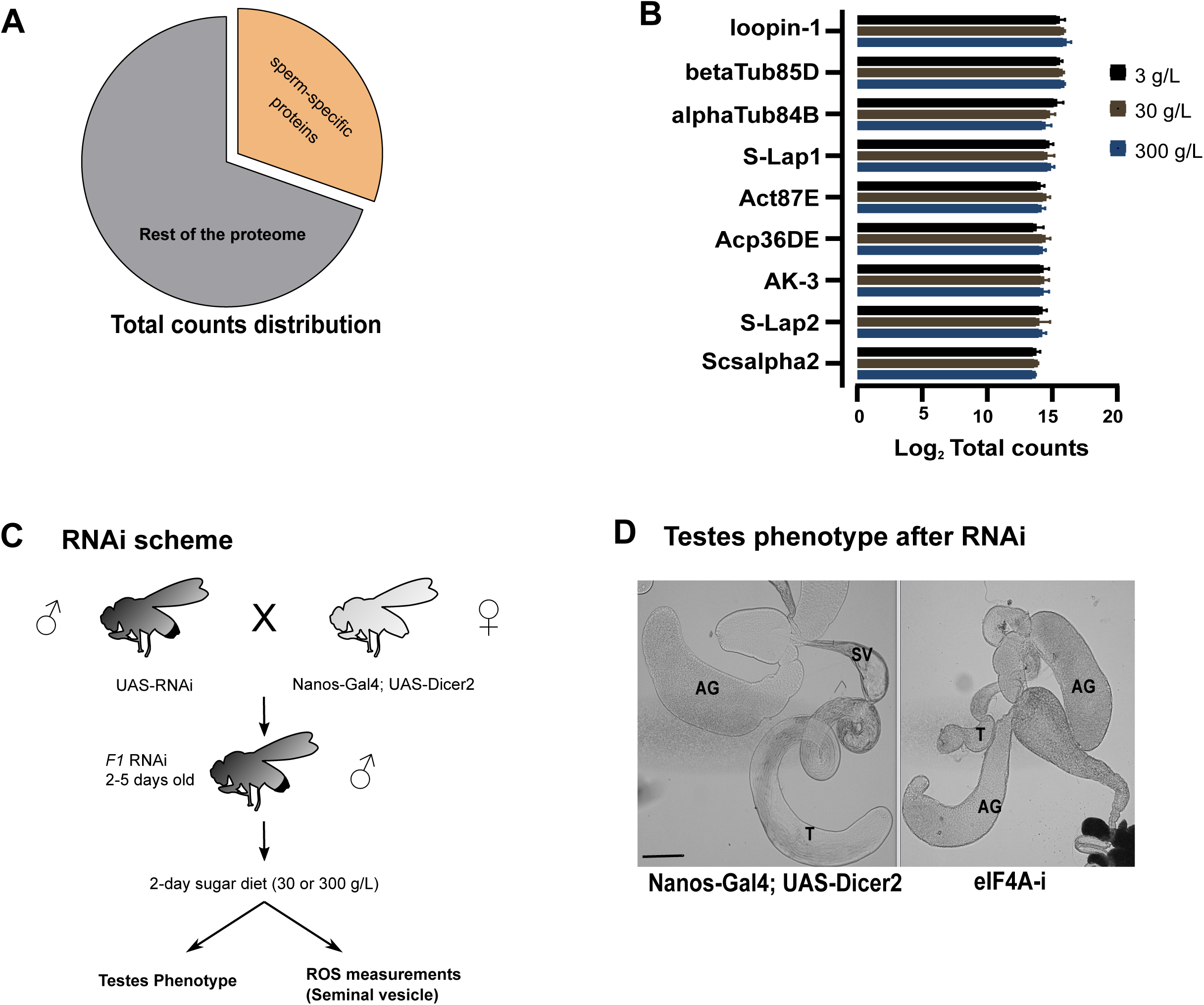
Proteomic composition of *Drosophila* sperm. **A:** The nine most abundant proteins are sperm specific and make up a third of the total counts. These sperm specific abundant proteins are indicated in orange within the pie-chart. Total counts distribution is presented. **B:** The expression of the top 9 abundant sperm proteins from **A** is not affected by dietary sugar as shown in the bar graph. Mean Log_2_ values are plotted in the bar graph. **C:** Schematics of RNAi crossing scheme. RNAi was induced in the germline by crossing virgin female Nanos-Gal4; UAS-Dicer2 flies with males carrying the UAS-RNAi construct. The F1 adults were screened for testes phenotype, and subjected to ROS measurements in seminal vesicles using ROS indicators **D:** Light-microscopy image of testes phenotype with and without eIF4A in the germline. DIC images of Nanos Gal4; Dicer2 heterozygous testes shows normal testes (T), accessory glands (AG) and seminal vesicles with sperm (SV). Nanos-Gal4; Dicer2> eIF4A RNAi (eIF4Ai) shows shorter T and lack SV. Scale bar: 700 µm DIC= differential interference contrast *Related to Figure 1*

**Figure S2:**
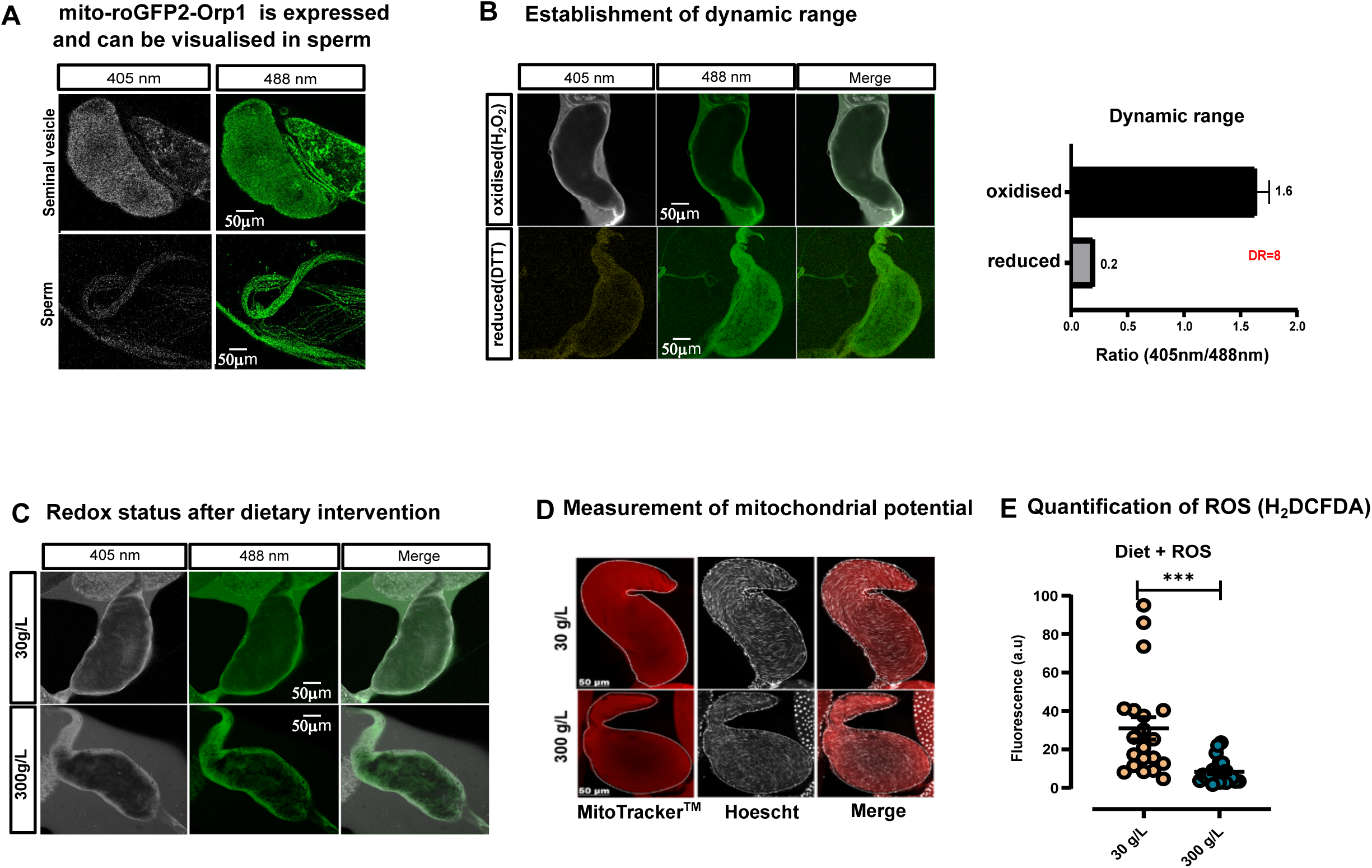
**A:** The mito-roGFP2-Orp1 is expressed and visualised in seminal vesicles (top panel) and on sperm (bottom panel). The sequential excitation wavelengths are 405- and 488 nm as indicated, and emission was captured at 518-580 nm. Scale bar: 50 µm **B:** Establishment of dynamic range for measurement of redox changes using the mito-roGFP2-Orp1 sensor. H_2_O_2_ and DTT represent fully oxidised and fully reduced conditions, respectively. Bar graphs represent quantified ratio of fluorescence emission from sequential excitation at 405-and 488 nm. Reduced condition was arbitrarily set at 0.2. DR= dynamic range. Scale bar: 50 µm **C:** Live imaging of roGFP2-Orp1 in sperm/seminal vesicles after dietary intervention with 30- or 300 g/L sugar. Scale bar: 50 µm **D:** MitoTracker™ Red-CMRos staining reveals no apparent change in mitochondrial potential (ΔΨm) in sperm/ seminal vesicle. Flies were fed 30- or 300 g/L sugar diet for two days, seminal vesciles were dissected, stained with the dye, and subjected to paraformaldehyde fixation. Fluorescence microscopy images showing cellular distribution of mitochondria-specific dye MitoTracker™ Red (red) in the sperm within the seminal vesicle. The tissues were counter stained for nuclei using Hoescht stain (White) and representative images for both diets are presented. Scale bar: 50µm **E:** Quantification of ROS labelled with H_2_DCFDA: The same flies in **D** were dissected, and seminal vesicles were immediately incubated with the ROS labelling dye-H_2_DCFDA and imaged. The micrographs were quantified and fluorescence (a.u) are plotted. The data shown are mean ± SEM. n=20, asterisks (*) represent p ≤ 0.001, unpaired t-test. *Related to Figure 2*

**Figure S3:**
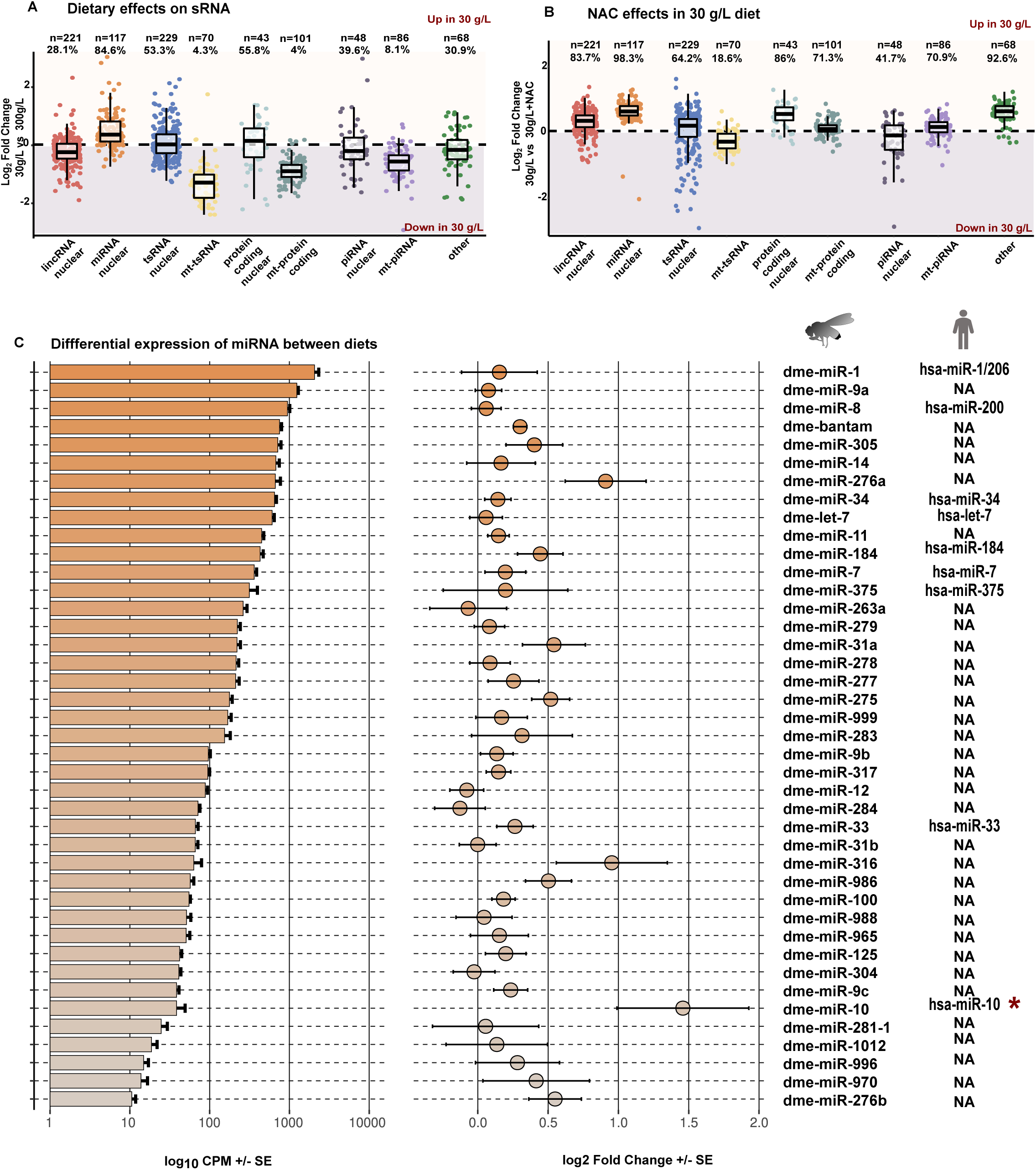
Biotype distribution in sperm is changed by dietary intervention. **A:** Fold changes 30-vs 300 g/L of the indicated sRNA biotypes are presented. Each dot represents a unique transcript. The reads mapping to nuclear and mitochondrial genome are separated. **B:** Fold changes 30 g/L vs 30 g/L with NAC of the indicated sRNA biotypes are presented. Each dot represents a unique transcript. The reads mapping to nuclear and mitochondrial genome are separated. **C:** Dietary change of miRNA in 30 g/L sugar vs 300 g/L. All miRNA identified in the study in fly sperm are presented under the fly pictogram. The corresponding human orthologues, where present, are indicated under the human pictogram. Bar graphs in descending order represent abundance (Log_10_CPM) of each miRNA. Each miRNA represents a mean of all transcripts annotated to that miRNA. Fold changes between 30-vs 300 g/L are indicated as closed circles. Asterisk (*) represents statistically significant miRNA (*p<0*.*05*) calculated by Generalized Linear Negative Binomial model, error bars are standard error (SE). *Related to Figure 3*

**Figure S4:**
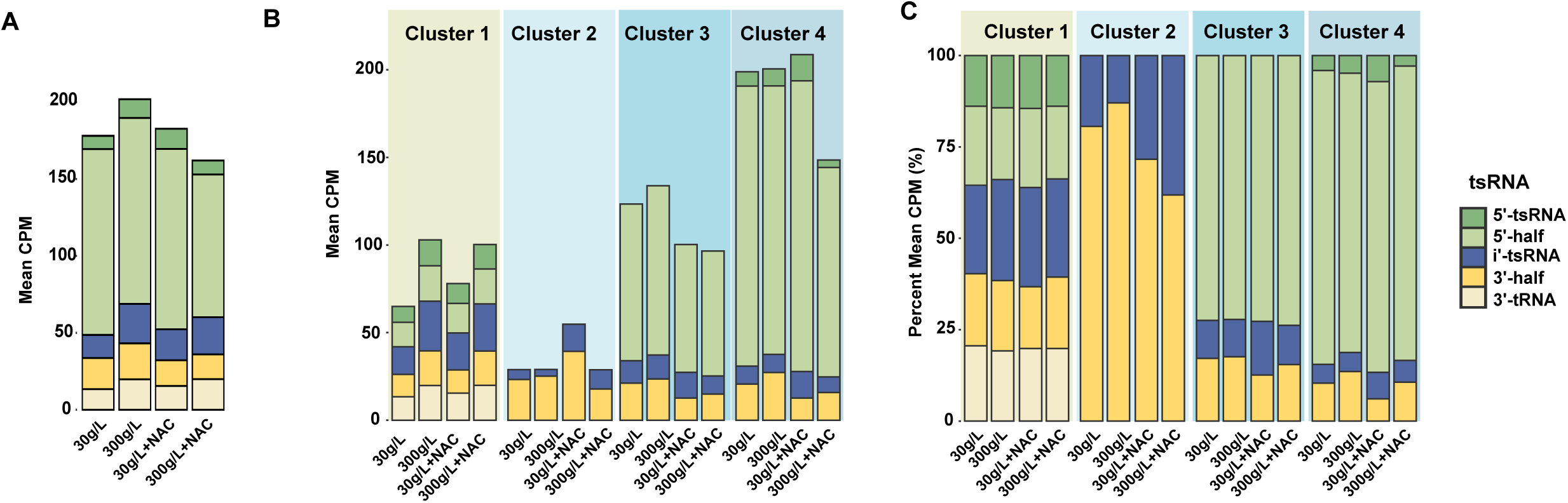
Distribution of tsRNA cleavage sites in cellular compartments. **A:** Mean CPM of all tsRNA, categorised on cleavage site. Each stacked bar represents one dietary condition. **B:** Mean CPM of tsRNA, categorised by cleavage site. Clusters 1-4 represent the hierarchical clustering as described for Figure 4C. **C:** Percentage of mean CPM of tsRNA, categorised by cleavage site. Stacked bars are divided by what hierarchical cluster they were classified as, according to heatmap in Figure 4C. *Related to Figure 4*

## Notes

### Competing Interest Statement

The authors have declared no competing interest.

